# The specificity of ParR binding determines the compatibility of conjugative plasmids in *Clostridium perfringens*

**DOI:** 10.1101/462192

**Authors:** Thomas D. Watts, Daouda A.K. Traore, Sarah C. Atkinson, Carmen Lao, Natalie Caltabiano, Julian I. Rood, Vicki Adams

**Affiliations:** Infection and Immunity Program, Monash Biomedicine Discovery Institute and Department of Microbiology, Monash University, Victoria 3800, Australia; Department of Biochemistry and Molecular Biology, Monash University, Victoria 3800, Australia; Faculté des Sciences et Techniques, Université des Sciences, des Techniques et des Technologies de Bamako (USTTB), BP E3206 Bamako, Mali

**Keywords:** Plasmid partitioning, Plasmid maintenance, Plasmid incompatibility, *Clostridium perfringens*, ParR, *parC*, surface plasmon resonance, DNA-binding, analytical ultracentrifugation

## Abstract

Plasmids that encode the same replication machinery are generally unable to coexist in the same bacterial cell. However, *Clostridium perfringens* strains often carry multiple conjugative toxin or antibiotic resistance plasmids that are closely related and encode similar Rep proteins. In many bacteria, plasmid partitioning upon cell division involves a ParMRC system and there are ~10 different ParMRC families in *C. perfringens*, with differences in amino acid sequences between each ParM family (15% − 54% identity). Since plasmids encoding genes belonging to the same ParMRC family are not observed in the same strain, these families appear to represent the basis for plasmid compatibility in *C. perfringens*. To understand this process, we examined the key recognition steps between ParR DNA-binding proteins and their *parC* binding sites. The ParR proteins bound to sequences within a *parC* site from the same ParMRC family, but could not interact with a *parC* site from a different ParMRC family. These data provide evidence that compatibility of the conjugative toxin plasmids of *C. perfringens* is mediated by their *parMRC*-like partitioning systems. This process provides a selective advantage by enabling the host bacterium to maintain separate plasmids that encode toxins that are specific for different host targets.

## INTRODUCTION

Low-copy number plasmids usually require an active partitioning system to ensure that they are faithfully inherited by daughter cells upon cell division (1). Type II or ParMRC plasmid partitioning systems encode three components: *parC*, a plasmid-encoded centromere, ParM, an actin-like ATPase that forms filaments in the presence of ATP or GTP and ParR, a DNA-binding adaptor protein that binds to *parC* (2-6). ParMRC systems stabilise the inheritance of plasmids by positioning them on either side of the cell septum prior to cell division.

ParR proteins are typically ribbon-helix-helix proteins that bind direct repeats within *parC*, either as a dimer or a dimer of dimers (5, 7-10). The *parC* centromere usually consists of a series of direct repeats upstream of the *parM* gene, however, its precise genetic structure differs between plasmids. Binding of ParR acts to seed the formation of a higher order solenoid-shaped structure, termed the segrosome, where the DNA wraps around ParR leaving a core of ParM interaction sites (9,10). Polymerising ParM filaments then link the ParR-*parC* complexes of two sister plasmids and push them to either cell pole (2,3,11-13). The initial step, in which ParR recognises and interacts with *parC*, is important in determining partition specificity between plasmids.

Plasmid incompatibility generally occurs when two co-resident plasmids encode the same essential replication or partitioning machinery (14). Most studies to date have focused on the partition specificity and incompatibility mediated by Type I or ParABS partitioning systems (15) (16-18), there is only limited evidence that partition-mediated incompatibility can also be facilitated by ParMRC-like partitioning systems (19).

In this study, we focused on partition-mediated incompatibility in *Clostridium perfringens*, a Gram-positive pathogen. In humans and animals *C. perfringens* produces an extensive range of toxins, which it uses to cause diseases that range from mild food poisoning to often fatal infections such as clostridial myonecrosis, enteritis and enterotoxaemia (20). Most *C. perfringens* toxins are encoded on large, low-copy number, conjugative plasmids (21) that are similar to the tetracycline resistance plasmid, pCW3 (21-27). These plasmids have approximately 35 kb of sequence similarity that includes the *tcp* conjugation locus and genes involved in replication, regulation and stable plasmid maintenance (Figure 1) (22,23,27-29). Even though these plasmids have similar replication regions, including a highly conserved replication protein, *C. perfringens* strains frequently carry up to five discrete plasmids (23,30). This phenomenon is typified by the avian necrotic enteritis isolate EHE-NE18, which stably maintains three large, closely related conjugative plasmids with Rep proteins that have 98% amino acid (aa) sequence identity (23,30).

**Figure 1.**
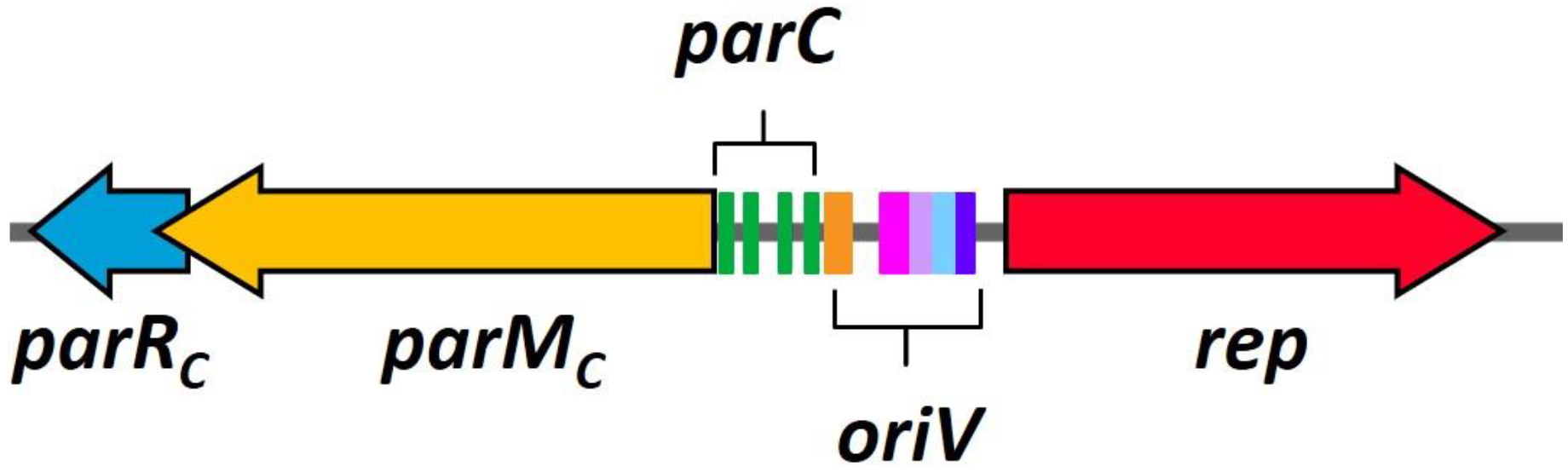
Replication and *parMRC* locus of pCW3: *parM*_C_ is shown in yellow, *parR*_C_ is shown in blue, the *parC*_C_ site (four direct repeats shown in green) is upstream of *parM*_C_, the five inverted repeats (IR) of *oriV* (IR1 in orange, IR2 in pink, IR3 in lavender, IR4 in blue, IR5 in purple) are shown upstream of *rep*, which is indicated by the red arrow.

Bioinformatic analysis has revealed the presence of at least ten families of ParMRC partitioning systems (ParMRC_A-J_) in these pCW3-like plasmids. The ParM components have > 90% aa sequence identity within a family, and 15-54% aa sequence identity between families, and the ParR and *parC* components show a similar trend (30). A representative of the ParMRC_B_ family was shown to be a true partitioning system, as addition of this partitioning system to an unstable mini-replicon was sufficient to stabilise its inheritance in *E. coli* (31). Strains of *C. perfringens* do not usually carry plasmids that encode the same ParMRC partitioning system (23,27,30), which suggests that these plasmids have evolved different partition specificities to ensure they are stably maintained within a single *C. perfringens* cell.

We recently showed that pCW3-like plasmids with identical partitioning systems could not be maintained in a single cell without selection, whereas plasmids with ParMRC systems from different families were stably maintained in *C. perfringens* cells (32). This finding suggested that differences in ParMRC plasmid partitioning systems were responsible for determining plasmid incompatibility between similar replicons and dictated which plasmid combinations could co-exist in an isolate. In this study, we have utilised Surface Plasmon Resonance (SPR) and Analytical Ultracentrifugation (AUC) to demonstrate that differences in the ParR and *parC* components of these partitioning families are reflected in their binding specificity, providing the essential biochemical evidence for the critical role of the ParMRC system in determining plasmid compatibility in *C. perfringens*.

## MATERIALS AND METHODS

### Plasmids, bacterial strains, and culture conditions

All *C. perfringens* strains, *Escherichia coli* strains and plasmids used in this study are listed in Table 1. All *E*. *coli* strains were grown on 2 × yeast tryptone (2YT) agar supplemented with 100 μg/ml of ampicillin and incubated at 37 °C overnight. *E. coli* expression strains were grown in either 2YT broth or autoinduction media (AIM) (33,34).

**Table 1.**
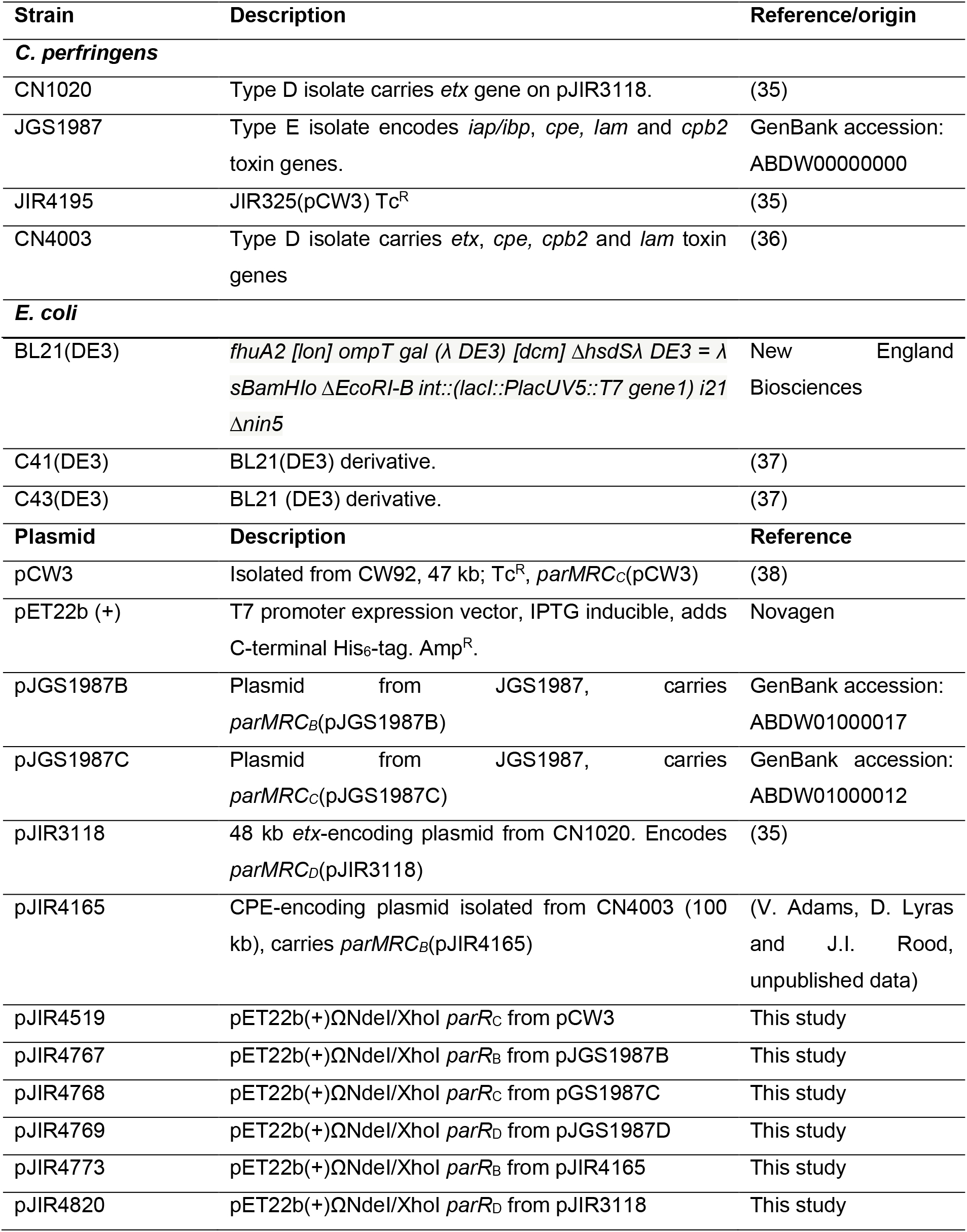
Bacterial strains and plasmids. TcR, tetracycline resistant, AmpR, ampicillin resistant. CPE, *C. perfringens* enterotoxin, *etx*, epsilon-toxin gene, lam, lamda toxin gene, *iap* and *ibp* iota-toxin genes, *cpb2*, beta2-toxin gene.

### Construction of ParR expression vectors

The *parR_C_* gene from pCW3 was codon optimised for expression in *E. coli*, synthesised by GenScript and cloned into the EcoRV site of pUC57-Kan. Codon optimised *parR_C_* then was subcloned into the NdeI/XhoI sites of pET22b(+). *parR_D_*(pJIR3118) was PCR amplified from CN1020 gDNA isolated as before (O’Connor *et al*., 2006) and cloned into the NdeI/Xhol site of pET22b (+) for expression. *parR_B_*(pJIR4165), *parR_B_*(pJGS1987B) *parR_C_*(pJGS1987C) and *parR_D_*(pJGS1987D) were codon optimised and synthesised before being cloned into pET22b(+) NdeI/XhoI sites by GenScript.

### ParR expression and purification

ParR proteins with C-terminal His6-tags were expressed using C43(DE3), C41(DE3) or BL21(DE3). *E. coli* cells were grown at either 28 °C in autoinduction media for 24 hours before lowering the temperature to 22 °C for 6 hours, or were grown in 2YT broth at 37 °C to an OD_600_ of 0.6 and induced with the addition of 0.1 mM IPTG for 3 hours (Supplementary Table 2). Cells were lysed using a cell disrupter (Avestin) (Lysis buffer: 20 mM TRIS (pH 7.9), 300 mM NaCl, 10% glycerol, 1 mg/ml DNase I and cOmplete protease inhibitors (Roche)) and proteins were purified (Supplementary figure 1) using TALON resin (Clontech) and eluted with the addition of increasing concentrations of imidazole (5 mM-200 mM) in purification buffer (20 mM TRIS (pH 7.9), 300 mM NaCl, 10% glycerol) and confirmed by Western blotting. All ParR proteins were buffer exchanged into buffer A (10 mM HEPES (pH 7.4), 300 mM NaCl, 3 mM EDTA, 0.05% Tween_20_, 0.02% NaN_3_) using a 3 kDa centrifugal filter (Amicon) before dilution to 0.1 μM. Independent preparations of each purified ParR protein were used as biological repeats for SPR.

### Fragment array preparation for SPR experiments

*parC* fragment arrays were constructed as previously described (39) using the Re-usable DNA capture technique (ReDCaT) method. Briefly, the *parC* regions of pCW3 (192 bp), pJIR3118 (230 bp) and pJIR4165 (262 bp) were used as templates for the Perl overlapping oligo program (POOP). POOP produced a series of overlapping forward and reverse 30 bp oligonucleotides (20 bp overlap). Reverse strand oligonucleotides had a 20 bp 3’ sequence (5’-CCTACCCTACGTCCTCCTGC-3’) that was complementary to the ReDCaT sequence (the ligands used in SPR experiments are listed in Supplementary Table 3). Oligonucleotides were synthesised (Integrated DNA technologies) at a concentration of 100 μM in IDTE buffer (10 mM Tris, 0.1 mM EDTA, pH 8.0). To construct fragments for SPR analysis, complimentary oligonucleotides were mixed in a ratio of 1.2:1 forward to reverse, annealed at 98 °C for 10 minutes and cooled for 30 minutes at room temperature. Fragments were then diluted to 0.5 nM in buffer A.

### Surface plasmon resonance

SPR experiments were based upon the ReDCaT method as previously described (39) and conducted using the Biacore T200 system (GE Healthcare Life Sciences). All experiments were carried out on an S series Biacore sensor chip (GE Healthcare Life Sciences) with streptavidin (SA) pre-immobilised to a carboxymethylated dextran matrix for capture of biotinylated interaction partners.

Prior to SPR, all four flow cells of the SA chip were washed three times with buffer containing 1 M NaCl and 50 mM NaOH. After washing and priming with buffer A, biotinylated ReDCaT linker (100 nM) (5’- biotin-GCAGGAGGACGTAGGGTAGG-3’) was immobilised to all four flow cells at 5 ul/min to a capture level of ~500 Response Units (RU). Subsequently, the chip was primed with buffer A and the ReDCaT complementary oligonucleotide (500 nM) was captured on flow cell 1, *parC* ligands diluted in buffer A to a concentration of 500 nM were captured to flow cells 2-4 (*parC*_B_, *parC*_C_, *parC*_D_ fragments on flow cells 2, 3 and 4 respectively) to a density of approximately 200 RU under flow conditions (10 μl/min for 30 seconds). DNA capture levels are listed in Supplementary Table 4. The first flow cell was used as a reference cell for subsequent measurements on flow cells 2 to 4. Each ParR protein (ParR_B_(pJIR4165), ParR_B_(pJGS1987B), ParR_C_(pCW3), ParR_C_(pJGS1987), ParR_D_(pJGS1987)) was diluted to a concentration of 0.1 μM in buffer A and ParR_D_(pJIR3118) was diluted in buffer A with 1 mg/ml dextran to reduce non-specific binding. Proteins were flowed through all four flow cells at 30 μl/min with 60 seconds association and 60 seconds dissociation. Binding stability measurements were recorded 10 seconds after the end of sample injection. All four flow cells of the chip were regenerated after each cycle using regeneration buffer (1 M NaCl and 50 mM NaOH) to leave only the biotinylated ReDCaT oligonucleotide. All experiments were conducted at 20 °C. All SPR methods were programmed using the Biacore T200 control software and data were analysed using the Biacore evaluation software version 2.0.

### Analytical Ultracentrifugation

Sedimentation velocity experiments were performed in an Optima Analytical Ultracentrifuge (Beckman Coulter) equipped with UV/Vis scanning optics. ParR_C_(pCW3) was prepared at a concentration of 0.5 mg/mL with and without 0.1 mg/mL *parC_C_* DNA (fragment C5). Reference (400 μL of buffer A without tween_20_) and sample (370 μL) solutions were loaded into double-sector cells with quartz windows. These cells were mounted in an An-50 Ti 8-hole rotor. Proteins and DNA were centrifuged at 40,000 rpm at 20°C, and radial absorbance data were collected at appropriate wavelengths (~280 nm) in continuous mode every 20 seconds. The partial specific volume (*v̄*) of ParR_C_ (0.7372), buffer density (1.0119 g/ml) and buffer viscosity (0.0104 P) was determined using the program SEDNTERP (40). The *v̄* of *parC_C_* C5 DNA (0.5500) was determined using UltraScan III (41). Data were fitted to continuous size-distribution [*c(s)*] and continuous mass distribution [*c*(*M*)] models using the program SEDFIT (42). All sedimentation coefficient data were normalised to standard conditions at 20°C in water (_*s*20,W_), relevant hydrodynamic properties are listed in Supplementary Table 5.

## RESULTS

### Identification of the pCW3 ParR_C_ binding site

The recognition steps between ParM, ParR and *parC* components both within and between different families of *parMRC* systems are likely to be key drivers in determining specificity of the partition reaction and therefore plasmid incompatibility in *C. perfringens*. The ParR-*parC* interaction is of particular interest as this is the first recognition step in the partitioning reaction (8,10,11) and is responsible for the incompatibility phenotype in some other plasmids (19).

Surface plasmon resonance (SPR) was employed to interrogate ParR-*parC* interactions. We first chose to examine the interaction between ParR_C_ and *parC_C_* from pCW3, as pCW3 is the most well characterised conjugative antimicrobial resistance plasmid in *C. perfringens* (28). To perform SPR, a recombinant His6-tagged ParR_C_(pCW3) protein was expressed in *E. coli* and purified (Supplementary Figure 1). A series of overlapping oligonucleotide fragments were designed (39) based on the 192 bp *parC_C_* region of pCW3 (Figure 1). These oligonucleotides were annealed to produce a fragment array consisting of 18 double-stranded *parC*_C_ fragments (denoted C1-C18) (Figure 2A). The stability and specificity of the ParR-*parC* interaction was assessed by challenging each *parC*C fragment with ParRC(pCW3) (Figure 2B + C). Strong interactions (a binding stability value >100 Response Units (RU)) between ParR_C_(pCW3) and fragments C1 (256 RU), C5 (249 RU), C6 (282 RU), C11 (154 RU), C12 (348 RU), C15 (217 RU) and C16 (311 RU) were observed. Weaker interactions (a stability value between baseline and 100 RU) were also noted for fragments C2 (54 RU), C7 (9 RU), C13 (48 RU) and C14 (42 RU). The strong interactions that were observed between *parC_C_*(pCW3) fragments and ParRC were mapped to the *parCC*(pCW3) nucleotide sequence, which showed that binding corresponded with the presence of four conserved 17 bp direct repeats (5’-AAACATCACAATTTTAC). The SPR results also indicate that a single fragment with the conserved *parCC* repeat was sufficient for ParR_C_ binding.

**Figure 2.**
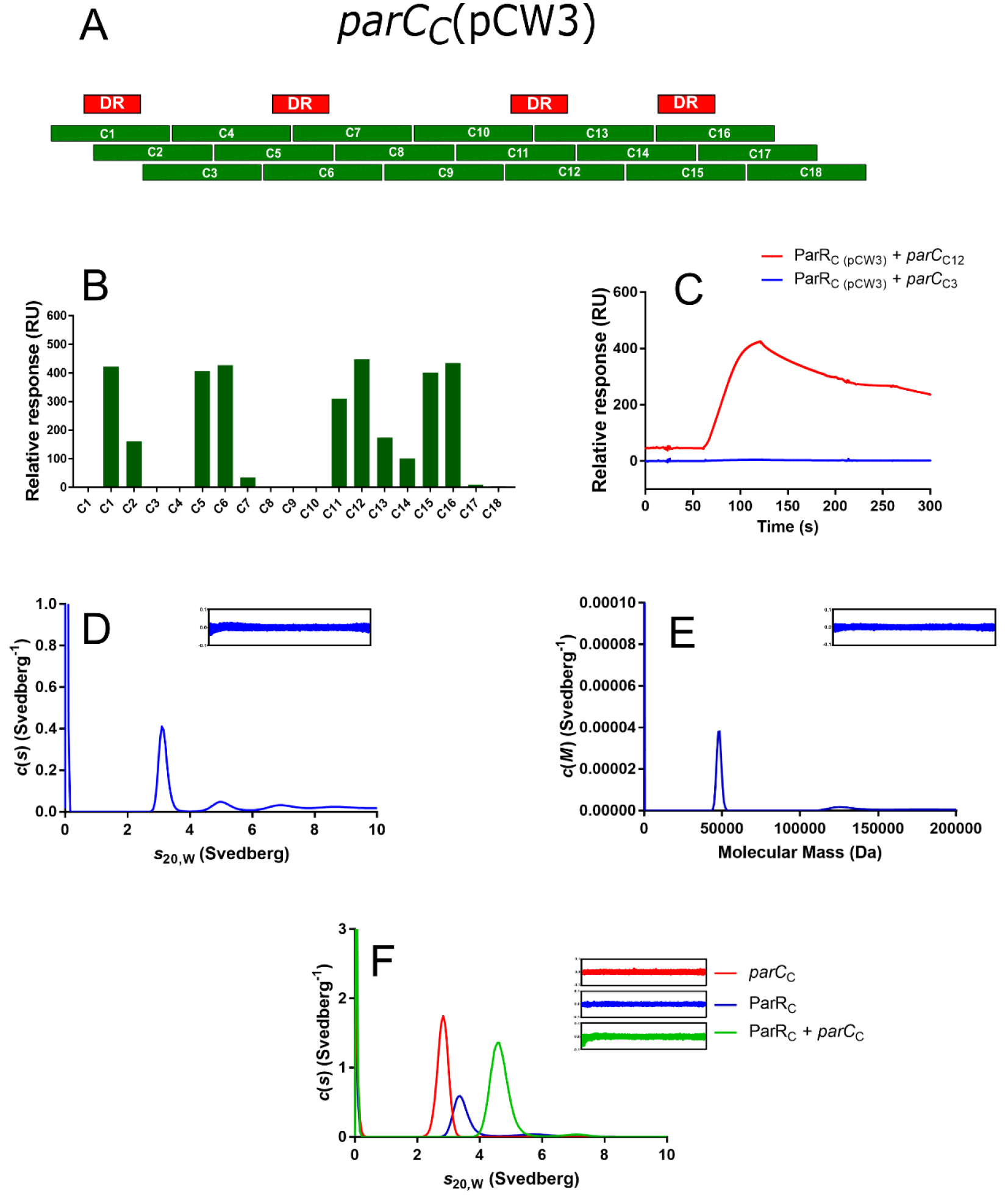
ParR_C_(pCW3) binds to a cognate *parC_C_*(pCW3) sequence. **(A)** Schematic of the *parC_C_*(pCW3) fragment array that consists of 30 bp fragments that overlap by 20 bp, direct repeats are indicated above the fragment array in red. **(B)** Representative ParR_C_(pCW3) binding to the *parC_C_*(pCW3) fragment array as determined by SPR. **(C)** Representative SPR binding curves for ParR_C_(pCW3) and *parC_C_*(pCW3) fragments, ParR_C_(pCW3) + C3 binding curve is shown in blue, and ParR_C_(pCW3) + C12 binding curve is shown in red. AUC sedimentation velocity experiments were also conducted on ParR_C_(pCW3), *parC_C_*(pCW3) fragment C5 and ParR_C_(pCW3) and *parC_C_*(pCW3) fragment C5 in combination. **(D)** The continuous sedimentation coefficient distribution [*c*(*s*)] as a function of normalised sedimentation coefficient (*s*_20,W_) for ParR_C_(pCW3). **(E)** The continuous mass distribution *c*(*M*) distribution as a function of molecular mass (Da) for ParR_C_(pCW3). **(F)** The continuous sedimentation coefficient distribution [*c*(*s*)] as a function of *s*_20,W_ for *parC_C_*(pCW3) C5 (red), ParR_C_(pCW3) (blue) and, ParR_C_(pCW3) and *parC_C_*(pCW3) C5 in combination (green). Residuals for each fit are shown as insets, confirming the validity of the fit of the data.

Analytical Ultracentrifugation (AUC) sedimentation velocity experiments were used to support the results obtained by SPR and provide insight into the multimeric state of ParRC in solution. The interaction between ParR_C_(pCW3) and the *parC_C_*(pCW3) fragment C5 was chosen for interrogation as the C5 fragment encodes a centrally located direct repeat and showed strong binding to ParR_C_(pCW3) by SPR. The results showed that ParR_C_(pCW3) primarily sedimented as a single species with a sedimentation coefficient (*s*_20,W_) of 3.1 S (Figure 2D), which corresponds to a molecular mass of 48 kDa (Figure 2E). The molecular mass of His_6_-tagged ParR_C_(pCW3) as predicted from the amino acid sequence is 10.9 kDa, suggesting that ParR_C_(pCW3) exists as a tetramer in solution. The *parC_C_*(pCW3) C5 fragment sedimented as a single species with a sedimentation coefficient of 2.7 S (Fig. 2F). When ParR_C_(pCW3) and *parC_C_*(pCW3) C5 were combined prior to centrifugation, a distinct shift in sedimentation coefficient to 4.2 S was observed (Figure 2F), which was consistent with binding in a 1:1 ratio of ParR_C_(pCW3) complex (four molecules) to each *parC*_C_(pCW3) binding site. This result confirmed that ParR_C_(pCW3) and *parC_C_*(pCW3) (C5) could interact in solution, which was consistent with the results obtained *via* SPR.

### ParR homologues cannot bind to non-cognate *parC* centromeres from a different phylogenetic ParMRC family

To determine if the interaction of ParR and *parC* components is ParMRC-family specific, two more ParR and *parC* families were included in the SPR analysis. ParR_B_ from pJIR4165 and ParR_D_ from pJIR3118 have 11% and 26% aa sequence identity to ParR_C_(pCW3), respectively, and were expressed and purified (Supplementary Figure 1). In addition, *parC*_B_(pJIR4165) and *parC*_D_(pJIR3118) fragment arrays were synthesised to yield fragments B1-B25 and D1-D21 (Figure 3A), these regions respectively have 45% and 47% nucleotide sequence identity to *parC*_C_(pCW3) (Supplementary Table 1). ParR_B_(pJIR4165), ParR_C_(pCW3) and ParR_D_(pJIR3118) were tested against each *parC* fragment array (*parC_B_*(pJIR4165)*, parC_C_*(pCW3) and *parC_D_*(pJIR3118)) in separate SPR experiments (Figure 3).

**Figure 3.**
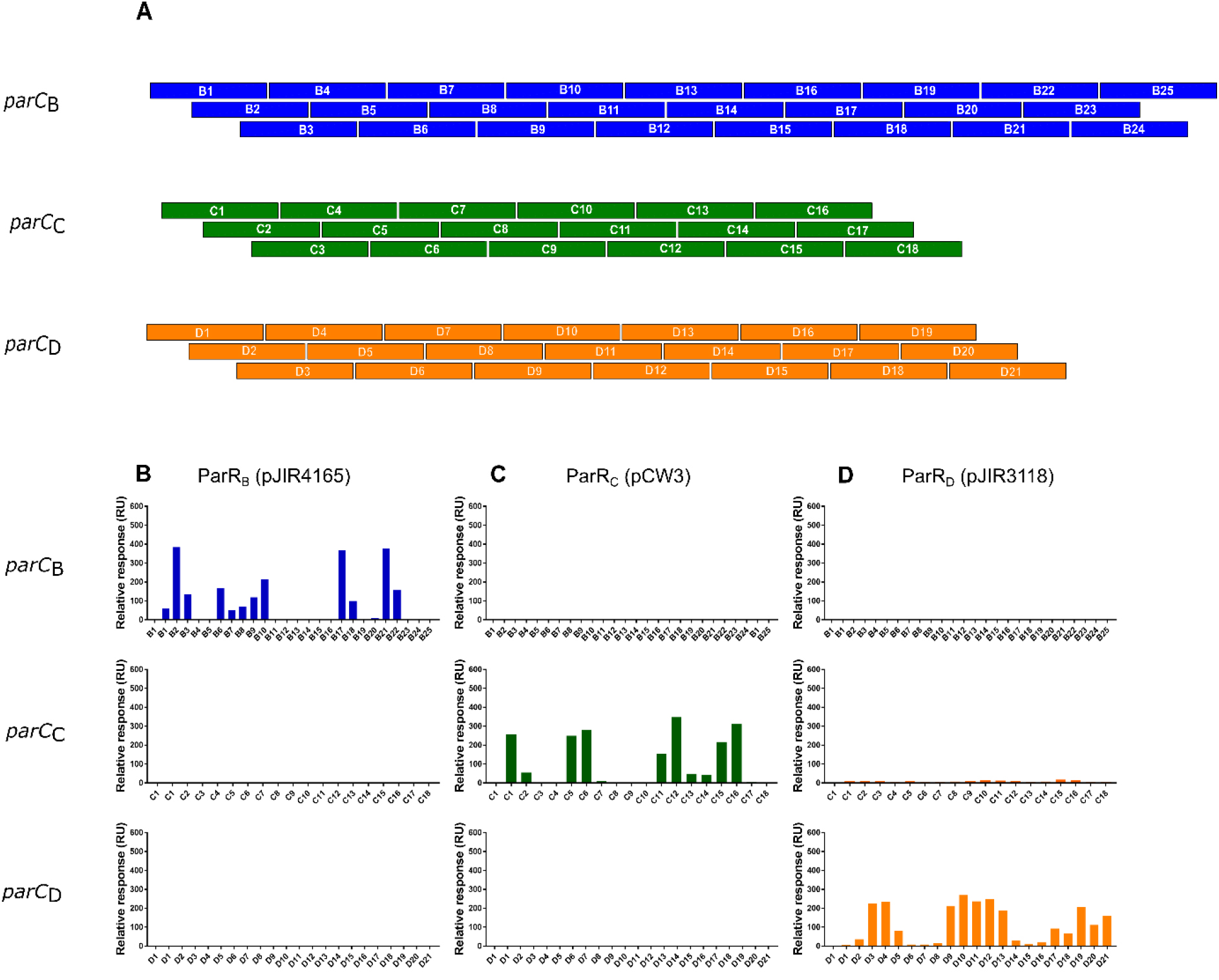
Surface plasmon resonance analysis demonstrated that ParR homologues bind to their cognate *parC* sites. A) Schematic of *parC* overlapping fragments. *parC_B_*(pJIR4165), *parC_C_*(pCW3) and *parC_D_*(pJIR3118) fragment arrays were constructed to test binding of ParR homologues to each *parC* region. All fragment arrays consisted of 30 bp oligonucleotides with 20 bp of overlapping sequence and were designed using POOP. Antisense oligonucleotides were constructed with the ReDCaT linker sequence present at the 3’ end of each fragment in the diagram above. Oligonucleotides were annealed before being captured onto the ReDCaT primed Streptavidin (SA) chip *via* the complementary base pairing between the ReDCaT linker and the complementary ReDCaT sequence on the Biacore T200 chip. B) SPR profiles obtained when ParR_B_(pJIR4165) was tested against *parC_B_*(pJIR4165) (blue), *parC_C_*(pCW3) (Green) and *parC_D_*(pJIR3118) (Orange) C) Shows ParR_C_(pCW3) binding profiles, D) Shows ParR_D_(pJIR3118) binding profiles. The first lane in every binding graph shows a no protein control with the fragments C1, B1 and D1.

The results showed that each ParR homologue bound only to its cognate *parC* fragment array. ParR_C_(pCW3) bound to its cognate *parC_C_* fragment array as before (Figure 3C), but did not bind non-cognate *parC_B_*(pJIR4165) or *parC_D_*(pJIR3118). ParR_B_(pJIR4165) bound to 12 *parC*_B_(pJIR4165) fragments with strongest binding (binding stability value of >300 RU) to fragments B2 (383 RU), B17 (368 RU) and B21 (377 RU) (Figure 3B). Unlike the *parC_C_*(pCW3) site, which had a clear correlation between binding and the direct repeat structures, the *parC*_B_(pJIR4165) region was more complex.

The *parC*_B_(pJIR4165) site consists of several different direct repeats and two inverted repeat structures, and many of these structures overlap. Therefore, mapping of ParR_B_(pJIR4165) binding to the *parC*_B_(pJIR4165) region did not indicate a clear ParR_B_(pJIR4165) binding site. ParR_C_(pCW3) was able to bind to its cognate *parC_C_*(pCW3) as before. ParR_C_(pCW3) was tested against *parC_B_*(pJIR4165) and *parC_D_*(pJIR3118) fragment arrays and showed no interaction with these non-cognate sequences (Figure 3C).

SPR analysis of the *parC_D_*(pJIR3118) fragment array with its cognate ParR_D_(pJIR3118) protein showed strong binding stability values (>100 RU) with fragments D3 (225 RU), D4 (232 RU), D9 (213 RU), D10 (270 RU), D11 (236 RU), D12 (250 RU) and D13 (187 RU) and weaker interactions (below 100 RU) with eight other oligonucleotide fragments (Figure 3D). Inspection of the *parC*_D_(pJIR3118) region revealed several different direct and inverted repeat structures, mostly consisting of variations of a conserved, AT-rich direct repeat (5’-TTATTTAAT).

However, mapping of the ParR_D_(pJIR3118) interactions did not give a clear indication of the specific ParR_D_ binding site. ParR_D_(pJIR3118) did not interact with the *parC_B_*(pJIR4165) fragment array and showed only very weak interactions with most of the fragments from the *parC*_C_(pCW3) array (stability values between 5-15 RU above baseline). These interactions are likely to be non-specific as a low level of binding was observed for all fragments, including the ReDCaT control fragment. The non-specific interactions were minimised by the addition of dextran to the SPR sample buffer, which had no effect on binding to the *parC_D_*(pJIR3118) fragments. Overall, these results highlight the specificity of the ParR-*parC* interactions, where ParR homologues only bind to their cognate *parC* component and have either no interaction or very weak inter-family interactions.

### ParR homologues recognise and bind non-cognate *parC* fragment arrays from the same ParRMC family

Our earlier work suggested that ParMRC components from the same family would be able to interact with one another, thus leading to interference with the partition process and plasmid incompatibility (32). To provide biochemical evidence for this hypothesis three different ParR homologues (ParR_B_, ParR_C_ and ParR_D_) from the *C. perfringens* strain JGS1987 were expressed, purified, (Supplementary Figure 1) and used to assess their capacity to facilitate intra-family interactions. There is an unpublished whole genome shotgun sequence available for strain JGS1987 (GenBank accession number: ABDW00000000) and it was chosen for analysis as an earlier bioinformatic survey revealed that this strain was particularly rich in *parMRC* genes (30). The JGS1987 sequence contains seven different *parM* alleles, which suggests that there may be seven potential plasmids present in this strain. Since these plasmid sequences had not been closed or given plasmid names, each putative plasmid was designated based on the strain of origin and the *parMRC* genes associated with that contig, yielding pJGS1987B, pJGS1987C and pJGS1987D etc. The JGS1987 ParR_B_, ParR_C_ and ParR_D_ homologues have 96%, 96% and 95% aa identity to the equivalent ParR_B_(pJIR4165), ParR_C_(pCW3) and ParR_D_(pJIR3118) proteins (Supplementary Table 1) (43). The corresponding JGS1987 *parC* regions also show high levels (82% to 91%) of nucleotide sequence similarity to the equivalent homologues (Supplementary Table 1). We postulated that the respective JGS1987-derived ParR proteins would cross-react with *parC* arrays from other members of the same ParMRC family. To examine this hypothesis, we tested the existing suite of *parC* fragment arrays with the purified ParR homologues from JGS1987.

The JGS1987 ParR homologues interacted with non-cognate *parC* fragment arrays from the same ParMRC family, but not with non-cognate *parC* fragments from different families (Figure 4). ParR_B_(pJGS1987B) interacted with *parC_B_*(pJIR4165) with a comparable binding pattern to ParR_B_(pJIR4165) (Figure 4A). Strong binding stability (>200 RU) scores were recorded for interactions between ParR_B_(pJGS1987B) and *parC_B_*(pJIR4165) fragments B1, B2, B3, B6, B8, B9, B10, B17, B18, B20, B21, B22 and B25. Weaker binding stability scores were seen for fragments B4, B7, B11, B16 and B23.

**Figure 4.**
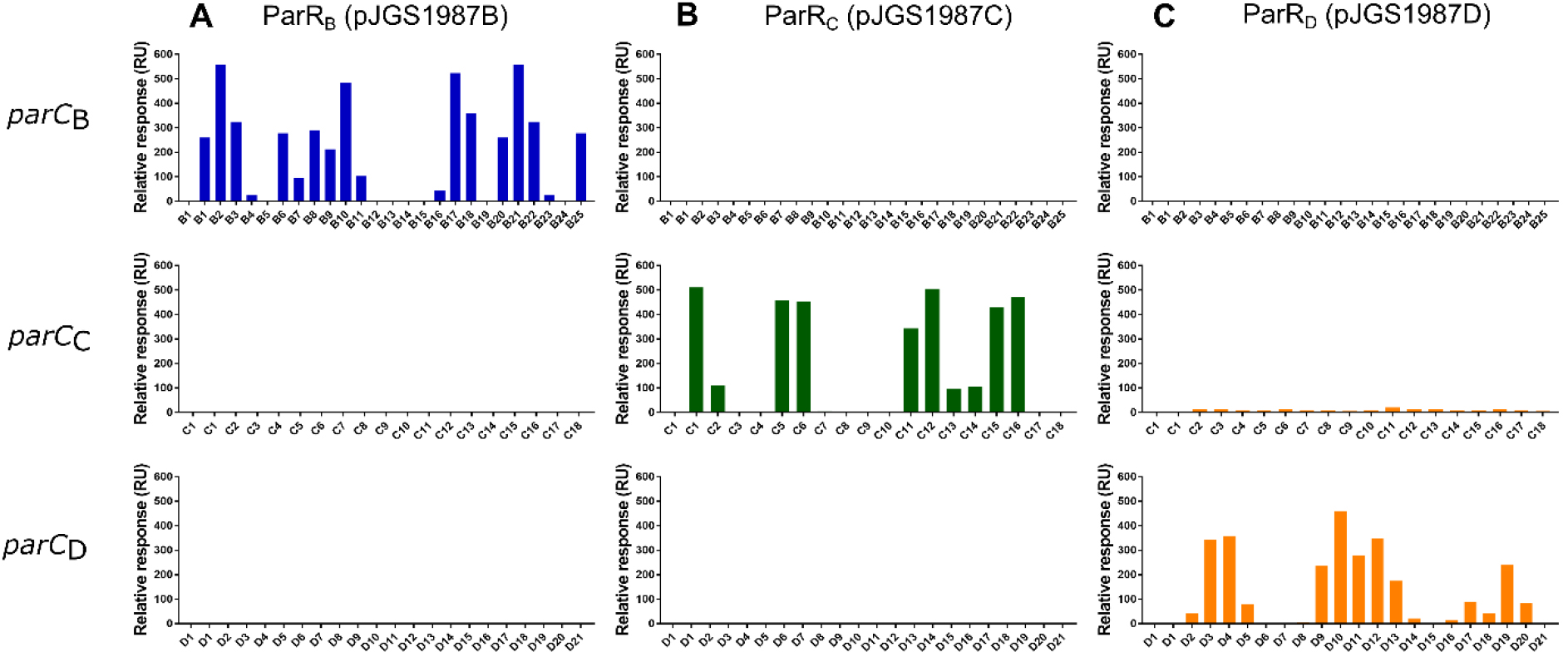
JGS1987 ParR homologues bind to non-cognate *parC* from the same family. ParR_B_, ParR_C_ and ParR_D_ homologues from the *C. perfringens* isolate, JGS1987, were tested against *parC_B_*(pJIR4165), *parC_C_*(pCW3) and *parC_D_*(pJIR3118) fragment arrays, and binding stability was measured using surface plasmon resonance. **A)** Shows ParR_B_(pJGS1987B) binding profiles when used to challenge *parC*_B_(pJIR4165) (blue), *parC_C_*(pCW3) and *parC_D_*(pJIR3118). **B)** Shows ParR_C_(pJGS1987C) binding profiles (blue) **C)** Shows ParR_D_(pJGS1987D) binding profiles (orange). The first fragment in every graph shows a no protein control

Similarly, ParR_C_(pJGS1987C) interacted only with *parC_C_*(pCW3), with the same binding pattern as observed for ParR_C_(pCW3) (Figure 4B). High binding stability (>200 RU) scores were recorded for interactions between ParRC(pJGS1987C) and *parC*_C_(pCW3) fragments C1, C5, C6, C11, C12 and C15. Weaker binding stability scores were recorded for C2, C13 and C14.

ParR_D_(pJGS1987D) only interacted with its non-cognate, but intra-family array from *parC_D_*(pJIR3118) (Figure 4C). Strong binding stability scores were recorded for interactions between ParR_D_(pJGS1987D) and *parC_D_*(pJIR3118) fragments D3, D4, D9, D10, D11, D12, D13 and D19. Whereas weaker binding stability scores were recorded for fragments D2, D5, D14, D16, D17, D18 and D20. Representative binding curves for each ParR-*parC* interaction pair are presented in Supplementary Figure 2. These data showed that ParR homologues interacted with non-cognate *parC* fragments from the same phylogenetic ParMRC family, thus confirming a subset of the bioinformatically derived phylogenetic groups of these homologues.

## DISCUSSION

In this study, we have demonstrated that ParR homologues from the pCW3-family of conjugative *C. perfringens* plasmids specifically recognise and bind to their cognate *parC* sites, providing biochemical evidence for the biological relevance of the phylogenetic ParMRC families that were previously identified (30). DNA binding studies showed that ParR proteins interacted with sequences within a centromeric *parC* site from the same ParMRC family, but could not interact with a non-cognate *parC* site from a different ParMRC family. We also demonstrated that ParR proteins can bind to non-cognate *parC* sites from the same ParMRC family (Figure 4). These findings are consistent with our previous phenotypic analysis of ParMRC-encoding plasmids in *C. perfringens*, where plasmids from the same partitioning family were unable to be maintained in a single *C. perfringens* isolate in the absence of selection (32). These combined data provide clear experimental evidence that variation in the ParMRC partitioning systems represents a major molecular mechanism by which native *C. perfringens* isolates can maintain multiple closely related plasmids in the same cell.

All ParR proteins characterised to date bind to directly repeated sequences, however, the repeats they interact with vary between plasmid systems. For example, ParR from the *E. coli* plasmid R1 requires a minimum of two 11 bp repeats for binding (11), ParR from pB171 (*E. coli*) binds two 10 bp direct repeats upstream of *parM* (44) and ParR from the *Staphylococcus aureus* plasmid pSK41 binds to 20 bp repeats (10).

The direct repeats in the *C. perfringens parC* sites differ substantially between families, with respect to both their nucleotide sequence and their spacing within the centromere. ParR_C_ binding correlated with four 17 bp direct repeats within the *parC*_C_ region. These repeat structures are conserved between *parC*_C_ regions of different plasmids, supporting the assertion that ParR is able to recognise and bind to these sites. By contrast, the ParR_B_ and ParR_D_ binding sites were more difficult to delineate because there were multiple direct and inverted repeat structures within the *parC*_B_ and *parC*_D_ regions.

Our findings support the hypothesis that the inability of ParR proteins to discriminate between closely related *parC* sites is responsible for previously observed ParMRC-mediated plasmid incompatibility (32). The consequence would be the incorrect linkage of two heterologous plasmids, eventually leaving distinct populations of daughter cells each containing only one of these plasmids (14,17,18,45). Although the heterologous pairing model is not favoured for type I partitioning mediated incompatibility (16,18), there is evidence that suggests this model could explain ParMRC-based plasmid incompatibility. For example, ParR from R1 is capable of linking replicons before partitioning and promiscuous binding of ParR from pB171 is responsible for plasmid incompatibility (8,19).

Analysis of our sedimentation velocity data showed that ParR_C_(pCW3) formed a tetrameric complex in solution. Upon the addition of a cognate *parC_C_* fragment containing the 17 bp direct repeat, a higher sedimentation coefficient was observed. This result provides additional evidence of the formation of specific complexes between ParR and *parC* recognition sequences in solution. These data are consistent with previous structural studies of ParR proteins from pSK41 and pB171 (9,10), which form tight dimers in solution and bind cooperatively to the DNA major groove within the *parC* centromere (5, 8-11). Once bound to *parC*, ParR forms a segrosome, where contacts between each ParR dimer are made, ultimately resulting in the formation of a dimer-of-dimers.

Replicon coevolution appears to be widespread in *C. perfringens*, where different isolates often carry closely related plasmids with different ParMRC partitioning systems (21,23,27). For example, the avian necrotic enteritis strain EHE-NE18 has three plasmids that have similar replication proteins, but different families of ParMRC system (ParMRC_A_, ParMRC_B_ and ParMRC_C_) (23). Based on the ParRB, ParRC and ParRD binding data reported here, and the previous genetic studies (32), it is concluded that to ensure that each plasmid is segregated independently these ParMRC systems have coevolved to carry different partition specificities.

The evolution of multiple ParMRC partition specificities in *C. perfringens* cells is reminiscent of the evolution of independent ParABS systems in *Burkholderia cenopacia*. The pathogenic *B. cenopacia* strain J2315 maintains three chromosomes and a large, low-copy number plasmid (46). The type I ParABS partitioning systems of these replicons have coevolved to become distinct so that each replicon is partitioned independently (46-49). Likewise, *Rhizobium leguminosarum bv. trifolii* RepB (ParB homologue) proteins discriminate between similar *parS* centromeres to independently segregate and maintain a chromosome in addition to four plasmids (50). Unlike *B. cenopacia* and *R. leguminosarum*, where the selection pressure to maintain multiple chromosomes and plasmids seems to have driven the coevolution of separate partition specificities, the selective pressure that has resulted in the generation of so many *parMRC* alleles in these conjugative *C. perfringens* plasmids remains unclear. One explanation may be that the ParMRC systems act as a means of competitive exclusion. It can be envisioned that upon entry into a new cell *via* conjugation, pCW3-like plasmids could displace resident plasmids that encode similar partitioning systems, thereby excluding them from the population. In addition, the plasmid-encoded toxin and antibiotic resistance genes may result in the positive selection of these plasmids in certain environmental niches, providing a selective advantage for the host cell if it can maintain these closely related plasmids. There is most certainly more complexity involved in the incompatibility phenotype in *C. perfringens*, since other factors such as the timing of plasmid replication, the plasmid copy number and plasmid replication initiation and regulatory proteins may play at least some role in determining whether two replicons are incompatible or are maintained in the same cell, as in other bacteria (14,15,18,51).

In conclusion, we have shown that interaction between the ParMRC partitioning components ParR and *parC* only occurs between members of the same phylogenetic family. These results provide biochemical insight into the basis of *C. perfringens* plasmid incompatibility and explain how multiple plasmids with similar replicons can be maintained within a single *C. perfringens* isolate.

## SUPPLEMENTARY DATA

Supplementary Data are available at NAR online.

## FUNDING

This research was supported by the Australian Research Council [grant DP160102680]. TDW was supported by the Australian Government Research Training Program (RTP). SCA was supported by a National Health and Medical Research Council Fellowship (1072267).

## REFERENCES

1. Gerdes, K., Howard, M. and Szardenings, F. (2010) Pushing and pulling in prokaryotic DNA segregation. Cell, 141, 927–942.

2. Moller-Jensen, J., Jensen, R.B., Lowe, J. and Gerdes, K. (2002) Prokaryotic DNA segregation by an actin-like filament. EMBO J., 21, 3119–3127.

3. van den Ent, F., Moller-Jensen, J., Amos, L.A., Gerdes, K. and Lowe, J. (2002) F-actin-like filaments formed by plasmid segregation protein ParM. EMBO J., 21, 6935–6943.

4. Breuner, A., Jensen, R.B., Dam, M., Pedersen, S. and Gerdes, K. (1996) The centromere-like *parC* locus of plasmid R1. Mol. Microbiol., 20, 581–592.

5. Dam, M. and Gerdes, K. (1994) Partitioning of plasmid R1. Ten direct repeats flanking the parA promoter constitute a centromere-like partition site *parC*, that expresses incompatibility. J. Mol. Biol., 236, 1289–1298.

6. Gerdes, K., Rasmussen, P.B. and Molin, S. (1986) Unique type of plasmid maintenance function: postsegregational killing of plasmid-free cells. Proc Natl Acad Sci U S A, 83, 3116–3120.

7. Jensen, R.B., Dam, M. and Gerdes, K. (1994) Partitioning of plasmid R1. The parA operon is autoregulated by ParR and its transcription is highly stimulated by a downstream activating element. J. Mol. Biol., 236, 1299–1309.

8. Jensen, R.B., Lurz, R. and Gerdes, K. (1998) Mechanism of DNA segregation in prokaryotes: replicon pairing by *parC* of plasmid R1. Proc Natl Acad Sci U S A, 95, 8550–8555.

9. Moller-Jensen, J., Ringgaard, S., Mercogliano, C.P., Gerdes, K. and Lowe, J. (2007) Structural analysis of the ParR/parC plasmid partition complex. EMBO J., 26, 4413–4422.

10. Schumacher, M.A., Glover, T.C., Brzoska, A.J., Jensen, S.O., Dunham, T.D., Skurray, R.A. and Firth, N. (2007) Segrosome structure revealed by a complex of ParR with centromere DNA. Nature, 450, 1268–1271.

11. Moller-Jensen, J., Borch, J., Dam, M., Jensen, R.B., Roepstorff, P. and Gerdes, K. (2003) Bacterial mitosis: ParM of plasmid R1 moves plasmid DNA by an actin-like insertional polymerization mechanism. Mol. Cell, 12, 1477–1487.

12. Gayathri, P., Fujii, T., Moller-Jensen, J., van den Ent, F., Namba, K. and Lowe, J. (2012) A bipolar spindle of antiparallel ParM filaments drives bacterial plasmid segregation. Science, 338, 1334–1337.

13. Bharat, T.A., Murshudov, G.N., Sachse, C. and Lowe, J. (2015) Structures of actin-like ParM filaments show architecture of plasmid-segregating spindles. Nature, 523, 106–110.

14. Novick, R.P. (1987) Plasmid incompatibility. Microbiological Reviews, 51, 381–395.

15. Ebersbach, G., Sherratt, D.J. and Gerdes, K. (2005) Partition-associated incompatibility caused by random assortment of pure plasmid clusters. Mol. Microbiol., 56, 1430–1440.

16. Bouet, J.Y., Rech, J., Egloff, S., Biek, D.P. and Lane, D. (2005) Probing plasmid partition with centromere-based incompatibility. Mol. Microbiol., 55, 511–525.

17. Funnell, B.E. (2005) Partition-mediated plasmid pairing. Plasmid, 53, 119–125.

18. Bouet, J.Y., Nordstrom, K. and Lane, D. (2007) Plasmid partition and incompatibility--the focus shifts. Mol. Microbiol., 65, 1405–1414.

19. Hyland, E.M., Wallace, E.W. and Murray, A.W. (2014) A model for the evolution of biological specificity: a cross-reacting DNA-binding protein causes plasmid incompatibility. J. Bacteriol., 196, 3002–3011.

20. Uzal, F.A., Freedman, J.C., Shrestha, A., Theoret, J.R., Garcia, J., Awad, M.M., Adams, V., Moore, R.J., Rood, J.I. and McClane, B.A. (2014) Towards an understanding of the role of *Clostridium perfringens* toxins in human and animal disease. Future microbiology, 9, 361–377.

21. Li, J., Adams, V., Bannam, T.L., Miyamoto, K., Garcia, J.P., Uzal, F.A., Rood, J.I. and McClane, B.A. (2013) Toxin plasmids of *Clostridium perfringens*. Microbiol. Mol. Biol. Rev., 77, 208–233.

22. Bannam, T.L., Teng, W.L., Bulach, D., Lyras, D. and Rood, J.I. (2006) Functional identification of conjugation and replication regions of the tetracycline resistance plasmid pCW3 from *Clostridium perfringens*. J. Bacteriol., 188, 4942–4951.

23. Bannam, T.L., Yan, X.X., Harrison, P.F., Seemann, T., Keyburn, A.L., Stubenrauch, C., Weeramantri, L.H., Cheung, J.K., McClane, B.A., Boyce, J.D. et al. (2011) Necrotic enteritis-derived *Clostridium perfringens* strain with three closely related independently conjugative toxin and antibiotic resistance plasmids. mBio, 2, e00190–00111.

24. Han, X., Du, X.D., Southey, L., Bulach, D.M., Seemann, T., Yan, X.X., Bannam, T.L. and Rood, J.I. (2015) Functional analysis of a bacitracin resistance determinant located on ICE*Cp1*, a novel Tn*916*-like element from a conjugative plasmid in *Clostridium perfringens*. Antimicrob. Agents Chemother., 59, 6855–6865.

25. Miyamoto, K., Fisher, D.J., Li, J., Sayeed, S., Akimoto, S. and McClane, B.A. (2006) Complete sequencing and diversity analysis of the enterotoxin-encoding plasmids in *Clostridium perfringens* type A non-food-borne human gastrointestinal disease isolates. J. Bacteriol., 188, 1585–1598.

26. Miyamoto, K., Li, J., Sayeed, S., Akimoto, S. and McClane, B.A. (2008) Sequencing and diversity analyses reveal extensive similarities between some epsilon-toxin-encoding plasmids and the pCPF5603 *Clostridium perfringens* enterotoxin plasmid. J. Bacteriol., 190, 7178–7188.

27. Parreira, V.R., Costa, M., Eikmeyer, F., Blom, J. and Prescott, J.F. (2012) Sequence of two plasmids from *Clostridium perfringens* chicken necrotic enteritis isolates and comparison with *C. perfringens* conjugative plasmids. PloS One, 7, e49753.

28. Wisniewski, J.A. and Rood, J.I. (2017) The Tcp conjugation system of *Clostridium perfringens*. Plasmid, 91, 28–36.

29. Traore, D.A.K., Wisniewski, J.A., Flanigan, S.F., Conroy, P.J., Panjikar, S., Mok, Y.F., Lao, C., Griffin, M.D.W., Adams, V., Rood, J.I. et al. (2018) Crystal structure of TcpK in complex with *oriT* DNA of the antibiotic resistance plasmid pCW3. Nature communications, 9, 3732.

30. Adams, V., Watts, T.D., Bulach, D.M., Lyras, D. and Rood, J.I. (2015) Plasmid partitioning systems of conjugative plasmids from *Clostridium perfringens*. Plasmid, 80, 90–96.

31. Chen, S., Larsson, M., Robinson, R.C. and Chen, S.L. (2017) Direct and convenient measurement of plasmid stability in lab and clinical isolates of *E. coli*. Scientific reports, 7, 4788.

32. Watts, T.D., Johanesen, P.A., Lyras, D., Rood, J.I. and Adams, V. (2017) Evidence that compatibility of closely related replicons in *Clostridium perfringens* depends on linkage to *parMRC*-like partitioning systems of different subfamilies. Plasmid, 91, 68–75.

33. Studier, F.W. (2005) Protein production by auto-induction in high density shaking cultures. Protein Expr Purif, 41, 207–234.

34. Studier, F.W. (2014) Stable expression clones and auto-induction for protein production in *E. coli*. Methods in molecular biology (Clifton, N.J.), 1091, 17–32.

35. Hughes, M.L., Poon, R., Adams, V., Sayeed, S., Saputo, J., Uzal, F.A., McClane, B.A. and Rood, J.I. (2007) Epsilon-toxin plasmids of *Clostridium perfringens* type D are conjugative. J. Bacteriol., 189, 7531–7538.

36. Sayeed, S., Fernandez-Miyakawa, M.E., Fisher, D.J., Adams, V., Poon, R., Rood, J.I., Uzal, F.A. and McClane, B.A. (2005) Epsilon-toxin is required for most *Clostridium perfringens* type D vegetative culture supernatants to cause lethality in the mouse intravenous injection model. Infect. Immun., 73, 7413–7421.

37. Miroux, B. and Walker, J.E. (1996) Over-production of proteins in *Escherichia coli*: mutant hosts that allow synthesis of some membrane proteins and globular proteins at high levels. J. Mol. Biol., 260, 289–298.

38. Rood, J.I., Scott, V.N. and Duncan, C.L. (1978) Identification of a transferable tetracycline resistance plasmid (pCW3) from *Clostridium perfringens*. Plasmid, 1, 563–570.

39. Stevenson, C.E., Assaad, A., Chandra, G., Le, T.B., Greive, S.J., Bibb, M.J. and Lawson, D.M. (2013) Investigation of DNA sequence recognition by a streptomycete MarR family transcriptional regulator through surface plasmon resonance and X-ray crystallography. Nucleic Acids Res., 41, 7009–7022.

40. Laue, T.M. (1992) Computer-aided interpretation of analytical sedimentation data for proteins. Analytical Ultracentrifugation in Biochemistry and Polymer Science, 90–125.

41. Demeler, B. (2005) In Scott, D. J., Harding, S. E. and Rowe, A. J. (eds.), Analytical Ultracentrifugation: Techniques and Methods. The Royal Society of Chemistry, (UK), pp. 210–230.

42. Schuck, P. (2000) Size-distribution analysis of macromolecules by sedimentation velocity ultracentrifugation and lamm equation modeling. Biophys. J., 78, 1606–1619.

43. McWilliam, H., Li, W., Uludag, M., Squizzato, S., Park, YM, Buso, N, Cowley, AP., Lopez, R.,. (2013) Analysis Tool Web Services from the EMBL-EBI. Nucleic Acids Res., 41, W597–600 doi:510.1093/nar/gkt1376.

44. Ringgaard, S., Ebersbach, G., Borch, J. and Gerdes, K. (2007) Regulatory cross-talk in the double par locus of plasmid pB171. J. Biol. Chem., 282, 3134–3145.

45. Austin, S. and Nordstrom, K. (1990) Partition-mediated incompatibility of bacterial plasmids. Cell, 60, 351–354.

46. Dubarry, N., Pasta, F. and Lane, D. (2006) ParABS systems of the four replicons of *Burkholderia cenocepacia*: new chromosome centromeres confer partition specificity. J. Bacteriol., 188, 1489–1496.

47. Passot, F.M., Calderon, V., Fichant, G., Lane, D. and Pasta, F. (2012) Centromere binding and evolution of chromosomal partition systems in the Burkholderiales. J. Bacteriol., 194, 3426–3436.

48. Du, W.L., Dubarry, N., Passot, F.M., Kamgoue, A., Murray, H., Lane, D. and Pasta, F. (2016) Orderly replication and segregation of the four replicons of *Burkholderia cenocepacia* J2315. PLoS Genet., 12, e1006172.

49. Pillet, F., Passot, F.M., Pasta, F., Anton Leberre, V. and Bouet, J.Y. (2017) Analysis of ParB-centromere interactions by multiplex SPR imaging reveals specific patterns for binding ParB in six centromeres of Burkholderiales chromosomes and plasmids. PloS One, 12, e0177056.

50. Koper, P., Zebracki, K., Marczak, M., Skorupska, A. and Mazur, A. (2016) RepB proteins of the multipartite *Rhizobium leguminosarum* bv. *trifolii* genome discriminate between centromere-like *parS* sequences for plasmid segregational stability. Mol. Microbiol., 102, 446–466.

51. Diaz, R., Rech, J. and Bouet, J.Y. (2015) Imaging centromere-based incompatibilities: Insights into the mechanism of incompatibility mediated by low-copy number plasmids. Plasmid, 80, 54–62.

